# Corruption of the Pearson correlation coefficient by measurement error: estimation, bias, and correction under different error models

**DOI:** 10.1101/671693

**Authors:** Edoardo Saccenti, Margriet H. W. B. Hendriks, Age K. Smilde

**Author notes:** These authors contributed equally to this work (Edoardo Saccenti, Age K. Smilde).

## Abstract

Correlation coefficients are abundantly used in the life sciences. Their use can be limited to simple exploratory analysis or to construct association networks for visualization but they are also basic ingredients for sophisticated multivariate data analysis methods. It is therefore important to have reliable estimates for correlation coefficients. In modern life sciences, comprehensive measurement techniques are used to measure metabolites, proteins, gene-expressions and other types of data. All these measurement techniques have errors. Whereas in the old days, with simple measurements, the errors were also simple, that is not the case anymore. Errors are heterogeneous, non-constant and not independent. This hampers the quality of the estimated correlation coefficients seriously. We will discuss the different types of errors as present in modern comprehensive life science data and show with theory, simulations and real-life data how these affect the correlation coefficients. We will briefly discuss ways to improve the estimation of such coefficients.

## 1 Introduction

The concept of correlation and correlation coefficient dates back to Bravais^1^ and Galton^2^ and found its modern formulation in the work of Fisher and Pearson^3, 4^, whose product moment correlation coefficient ***ρ*** has become the most used measure to describe the linear dependence between two random variables. From the pioneering work of Galton on heredity, the use of correlation (or co-relation as is it was termed) spread virtually in all fields of research and results based on it pervade the scientific literature.

Correlations are generally used to quantify, visualize and interpret bivariate (linear) relationships among measured variables. They are the building blocks of virtually all multivariate methods such as Principal Component Analysis (PCA^5–7^), Partial Least Squares regression, Canonical Correlation Analysis (CCA^8^) which are used to reduce, analyze and interpret high-dimensional *omics* data sets and are often the starting point for the inference of biological networks such as metabolite-metabolite associations networks^9, 10^, gene regulatory networks^11, 12^ an co-expression networks^13, 14^.

Fundamentally, correlation and correlation analysis are pivotal for understanding biological systems and the physical world. With the increase of comprehensive measurements (liquid-chromatography mass-spectrometry, nuclear magnetic resonance, gas-chromatography mass-spectrometry in metabolomics and proteomics; RNA-sequencing in transcriptomics) in life sciences, correlations are used as a first tool for visualization and interpretation, possibly after selection of a threshold to filter the correlations. However, the complexity and the difficulty of estimating correlation coefficients is not fully acknowledged.

Measurement error is intrinsic to every experimental technique and measurement platform, be it a simple ruler, a gene sequencer or a complicated array of detectors in a high-energy physics experiment, and in the early days of statistics it was known that measurement errors can bias the estimation of correlations^15^. This bias was called attenuation because it was found that under the error condition considered, the correlation was attenuated towards zero. The attenuation bias has been known and discussed in some research fields^16–19^ but it seems to be totally neglected in modern *omics*-based science. Moreover, contemporary comprehensive *omics* measurement techniques have far more complex measurement error structures than the simple ones considered in the past and on which early results were based.

In this paper, we intend to show the impact of measurement errors on the quality of the calculated correlation coefficients and we do this for several reasons. First, to make the *omics* community aware of the problems. Secondly, to make the theory of correlation up to date with current *omics* measurements taking into account more realistic measurement error models in the calculation of the correlation coefficient and third, to propose ways to alleviate the problem of distortion in the estimation of correlation induced by measurement error. We will do this by deriving analytical expressions supported by simulations and simple illustrations. We will also use real-life metabolomics data to illustrate our findings.

## 2 Measurement error models

We start with the simple case of having two correlated biological entities *x*_0_ and *y*_0_ which are randomly varying in a population. This may, e.g., be concentrations of two blood-metabolites in a cohort of persons or gene-expressions of two genes in cancer tissues. We will assume that those variables are normally distributed

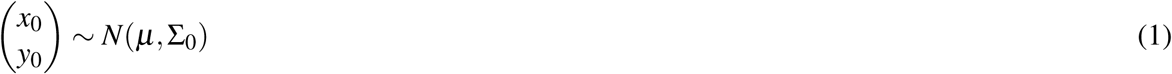

with underlying parameters

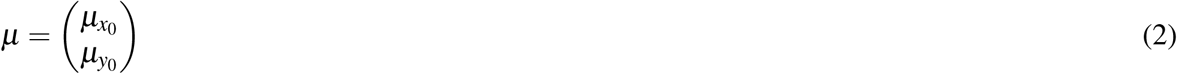

and

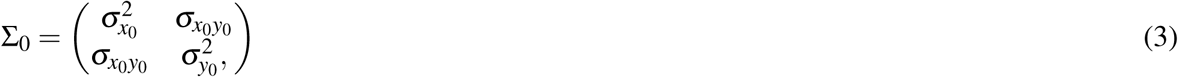

Under this model the variance components 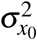 and 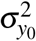 describe the biological variability for *x*_0_ and *y*_0_, respectively. The correlation ***ρ***_0_, between *x*_0_ and *y*_0_ is given by

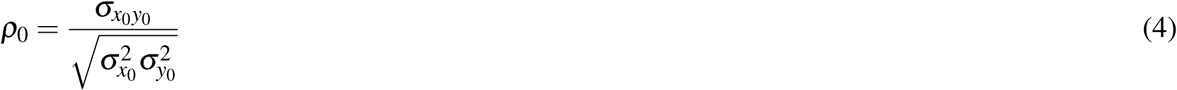

We refer to ***ρ***_0_ as the *true correlation*.

Whatever the nature of the variables *x*_0_ and *y*_0_ and whatever the experimental technique used to measure them there is always a random error component (also refereed to as noise or uncertainty) associated with the measurement procedure. This random error is by its own nature not reproducible (in contrast with systematic error which is reproducible and can be corrected for) but can be modeled, *i.e.* described, in a statistical fashion. Such models have been developed and applied in virtually every area of science and technology and can be used to adjust for measurement errors or to describe the bias introduced by it. The measured variables will be indicated by *x* and *y* to distinguished them from *x*_0_ and *y*_0_ which are their errorless counterparts.

The correlation coefficient ***ρ***_0_ is sought to be estimated from these measured data. Assuming that *N* samples are taken, the sample correlation *r*_*N*_ is calculated as

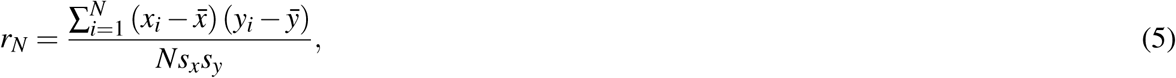

where 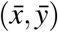 is the sample mean over *N* observations and *s*_*x*_, *s*_*y*_ are the usual sample standard deviation estimators. This sample correlation is used as a proxy of ***ρ***_0_. The population value of this sample correlation is

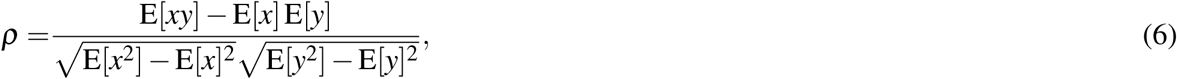

and it also holds that

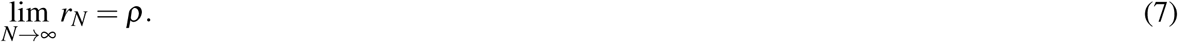

We will call ***ρ*** the *expected correlation*. Ideally, ***ρ***_0_ = ***ρ*** but this is unfortunately not always the case. In plain words: certain measurement errors do not cancel out if the number of samples increases.

In the following section we will introduce three error models and will show with both simulated and real data how measurement error impacts the estimation of the Pearson correlation coefficient. We will focus mainly on ***ρ***_0_ and ***ρ***.

### 2.1 Additive error

The most simple error model is the additive error model where the measured entities *x* and *y* are modeled as

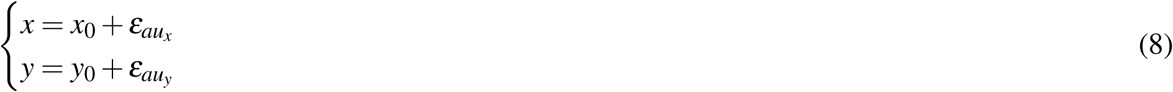

where it is assumed that the error components 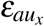 and 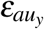 are independently normally distributed around zero with variance 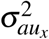 and 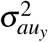 and are also independent from *x*_0_ and *y*_0_. The subscripts *au*_*x*_, *au*_*y*_ stand for *a*dditive *u*ncorrelated error (*ε*) on variable *x* and *y*.

Variable *x* and *y* represent measured quantities accessible to the experimenter. This error model describes the case in which the measurement error causes within-sample variability, which means that *p* measurement replicates *x*_*i*,1_, *x*_*i*,2_, … *x*_*i,p*_ of observation *x*_*i*_ of variable *x* will all have slightly different values due to the random fluctuation of the error component 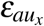; the extent of the variability among the replicates depends on the magnitude of the error variance 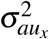 (and similarly for the *y* variable). This can be seen in Figure 1A where it is shown that in the presence of measurement error (*i.e.* 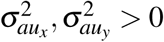) the two variables *x* and *y* are more dispersed. Due to the measurement error, the expected correlation coefficient ***ρ*** is always biased downwards, *i.e. **ρ*** < ***ρ***_0_, as already shown by Spearman^15^ (see Figure 1B) who also provided an analytical expression for the attenuation of the expected correlation coefficient as a function of the error components (a modern treatment can be found in reference^20^):

**Figure 1.**
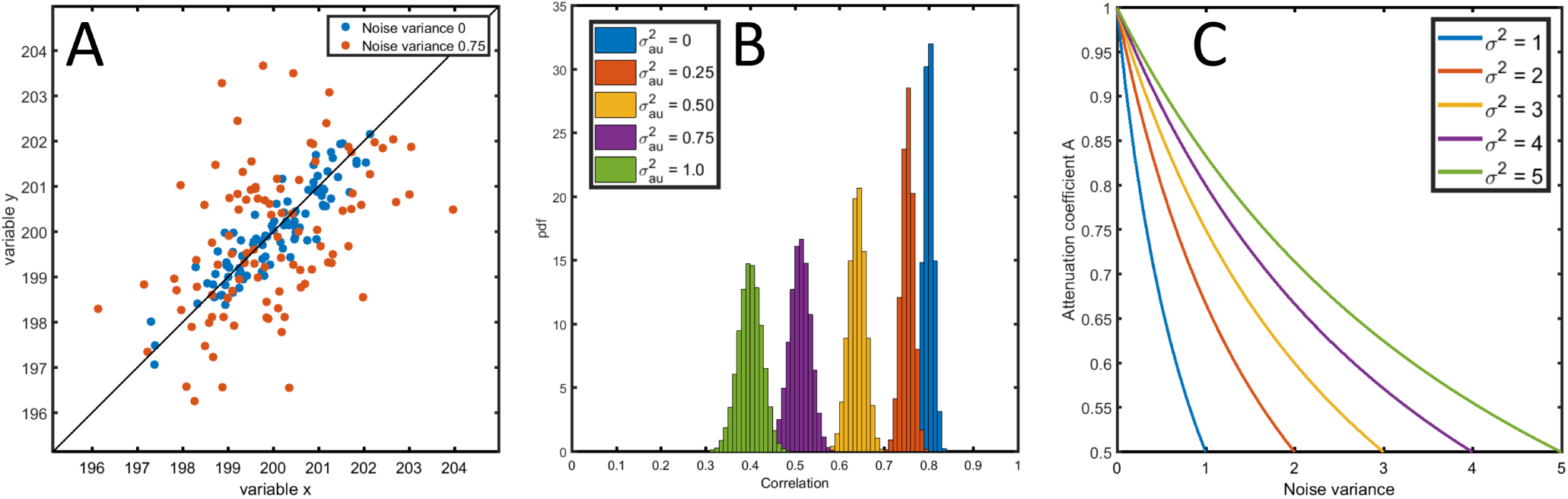
**A**: Correlation plot of two variables *x* and 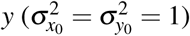 generated without 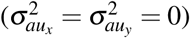 and with uncorrelated additive error 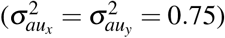 with underlying true correlation ***ρ***_0_ = 0.8 (model 8). **B**: Distribution of the sample correlation coefficient for different levels of measurement error 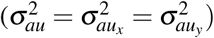 for a true correlation ***ρ***_0_ = 0.8.**C**: The attenuation coefficient *A* from Equation (10) as a function the measurement error for different level of the variance 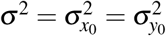 of the variables *x*_0_ and *y*_0_. See Material and Methods section 6.5.1 for details on the simulations.

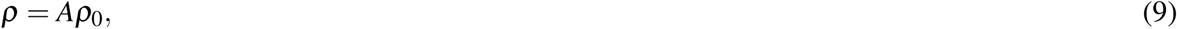

where

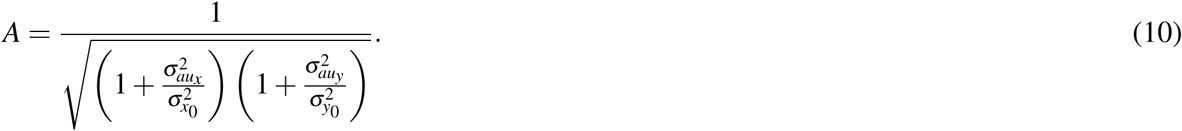

Equation (9) implies that in presence of measurement error the expected correlation is different from the true correlation ***ρ***_0_ which is sought to be estimated. The attenuation *A* is always strictly smaller than 1 and it is a decreasing function of the size of the measurement error relative to the biological variation (see Figure 1C), as it can be seen from Equation (10). The attenuation of the expected correlation, despite being known since 1904, has sporadically resurfaced in the statistical literature in the psychological, epidemiology and behavioral sciences (where it is known as attenuation due to intra-person or intra-individual variability, see^19^ and reference therein) but has been largely neglected in the life sciences, despite its relevance.

The error model (8) can be extended to include a correlated error term *ε*_*ac*_

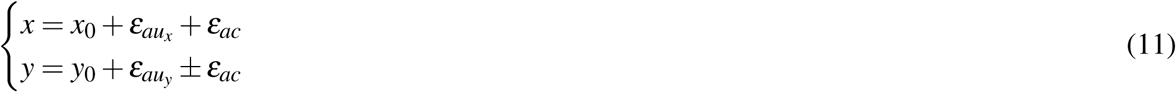

with *ε*_*ac*_ normally distributed around zero with variance 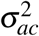. The ‘±’ models the sign of the error correlation. When *ε*_*ac*_ has a positive sign in both *x* and *y* the error is positively correlated; if the sign is discordant the error is negatively correlated. The subscript *ac* is used to indicate *a*dditive *c*orrelated error. The variance for *x* is given by

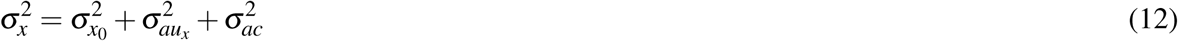

and likewise for the variable *y*. In general, additive correlated error can have different causes depending on the type of instruments and measurement protocols used. For example, in transcriptomics, metabolomics and proteomics, usually samples have to be pretreated (sample work-up) prior to the actual instrumental analysis. Any error in a sample work-up step may affect all measured entities in a similar way^21^. Another example is the use of internal standards for quantification: any error in the amount of internal standard added may also affect all measured entities in a similar way. Hence, in both cases this leads to (positively) correlated measurement error. In some cases in metabolomics and proteomics the data are preprocessed using deconvolution tools. In that case two co-eluting peaks are mathematically separated and quantified. Since the total area under the curve is constant and (positive) error in one of the deconvoluted peaks is compensated by a (negative) error in the second peak, this may give rise to negatively correlated measurement error.

To show the effect of additive uncorrelated measurement error we consider the concentration profiles of three hypothetical metabolites P1, P2 and P3 simulated using a simple dynamic model (see Figure 2A and Section 6.5.2) where additive uncorrelated measurement error is added before calculating the pairwise correlations among P1, P2 and P3: also in this case the magnitude of the correlation is attenuated, and the attenuation increases with the error variance (see Figure 2B.)

**Figure 2.**
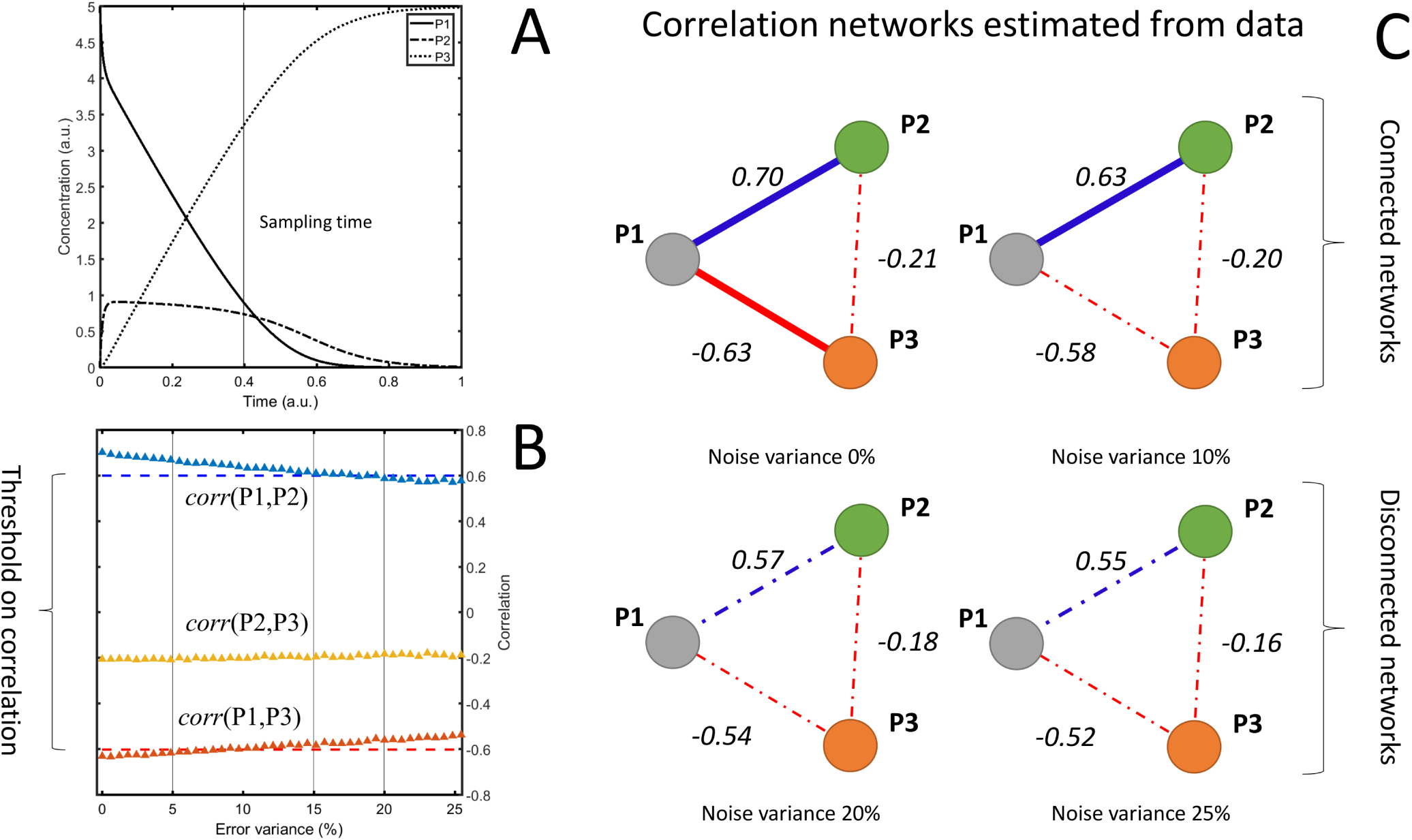
Consequences of measurement error when using correlation in systems biology. **A**: Time concentration profile of three metabolites P1, P2 and P3 generated through a simple enzymatic metabolic model; 100 profiles are generated by randomly varying the kinetic parameters defining the model and sampled at time 0.4 (a.u.). **B**: Average pairwise correlation of P1, P2 and P3 as a function of the variance of the additive uncorrelated error. **C**: Inference of a metabolite-metabolite correlation network: two metabolites are associated if their correlation is above 0.6^22^ (see threshold in **B**). The increasing level of measurement error hampers the network inference (compare the different panels). See Material and Methods section 6.5.2 for details on the simulations.

This has serious repercussions when correlations are used for the definition of association networks, as commonly done in systems biology and functional genomics^10, 23^: measurement error drives correlation towards zero and this impacts network reconstruction. If a threshold of 0.6 is imposed to discriminate between correlated and non correlated variables as usually done in metabolomics^22^, an error variance of around 15% (see Figure 2B, point where the correlation crosses the threshold) of the biological variation will attenuate the correlation to the point that metabolites will be deemed not to be associated even if they are biologically correlated leading to very different metabolite association networks (see Figure 2C.)

### 2.2 Multiplicative error

In many experimental situations it is observed that the measurement error is proportional to the magnitude of the measured signal; when this happens the measurement error is said to be multiplicative. The model for sampled variables in presence of multiplicative measurement error is

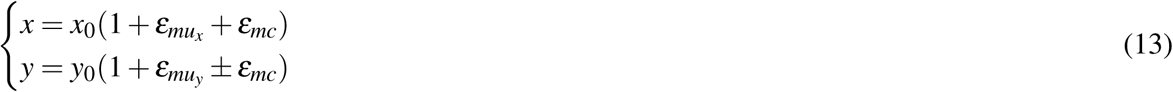

where 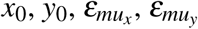 and *ε*_*mc*_ have the same distributional properties as before in the additive error case, and the last three terms represent the *m*ultiplicative *u*ncorrelated errors in *x* and *y*, respectively, and the *m*ultiplicative *c*orrelated error.

The characteristics of the multiplicative error and the variance of the measured entities 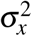 depend on the level *µ*_*x*0_ of the signal to be measured (for a derivation of Equation (14) see Section 6. 6.1.1):

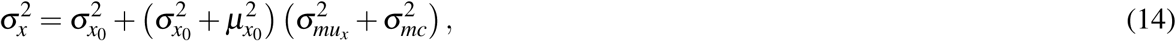

while in the additive case the standard deviation is similar for different concentrations and does not depend explicitly on the signal intensity, as shown in Equation (12). A similar equation holds for the variable *y*.

It has been observed that multiplicative errors often arises because of the different procedural steps like sample aliquoting^24^: this is the case of deep sequencing experiments where the multiplicative error is possibly introduced by the pre-processing steps like, for example, linker ligation and PCR amplification which may vary from tag to tag and from sample to sample^25^. In other cases the multiplicative error arise from the distributional properties of the signal, like in those experiments where the measurement comes down to counts like in the case of RNA fragments in an RNA-seq experiment or numbers of ions in a mass-spectrometer that are governed by Poisson distributions for which the standard deviation is equal to the mean. For another example, in NMR spectroscopy measured intensities are affected by the sample magnetization conditions: fluctuations in the external electromagnetic field or instability of the rf pulses affects the signal in a fashion that is proportional to the signal itself^26^.

A multiplicative error distorts correlations and this affects the results of any data analysis approach which is based on correlations. To show the effect of multiplicative error we consider the analysis of a simulated metabolomic data set starting from real mass-spectrometry (MS) data on which extra uncorrelated and correlated multiplicative measurement errors have been added. As can be seen in Figure 3A the addition of error affects the underlying data structure: the error free data is such that only a subset of the measured variables contributes to explain the pattern in a low dimensional projection of the data, *i.e.* have PCA loadings substantially different from zero (3B). The addition of extra multiplicative error perturbs the loading structure to the point that all variables contribute equally to the model (3C), obscuring the real data structure and hampering the interpretation of the PCA results. This is not necessarily caused by the multiplicative nature of the error, it is certainly caused by the correlated error part. Since the term *ε*_*mc*_ is common to all variables it introduces the same amount of correlation among all the variables and this leads to all the variables contributing to the latent vector (principal component). One may also observe that the variation explained by the first principal component increases when adding the correlated measurement error.

**Figure 3.**
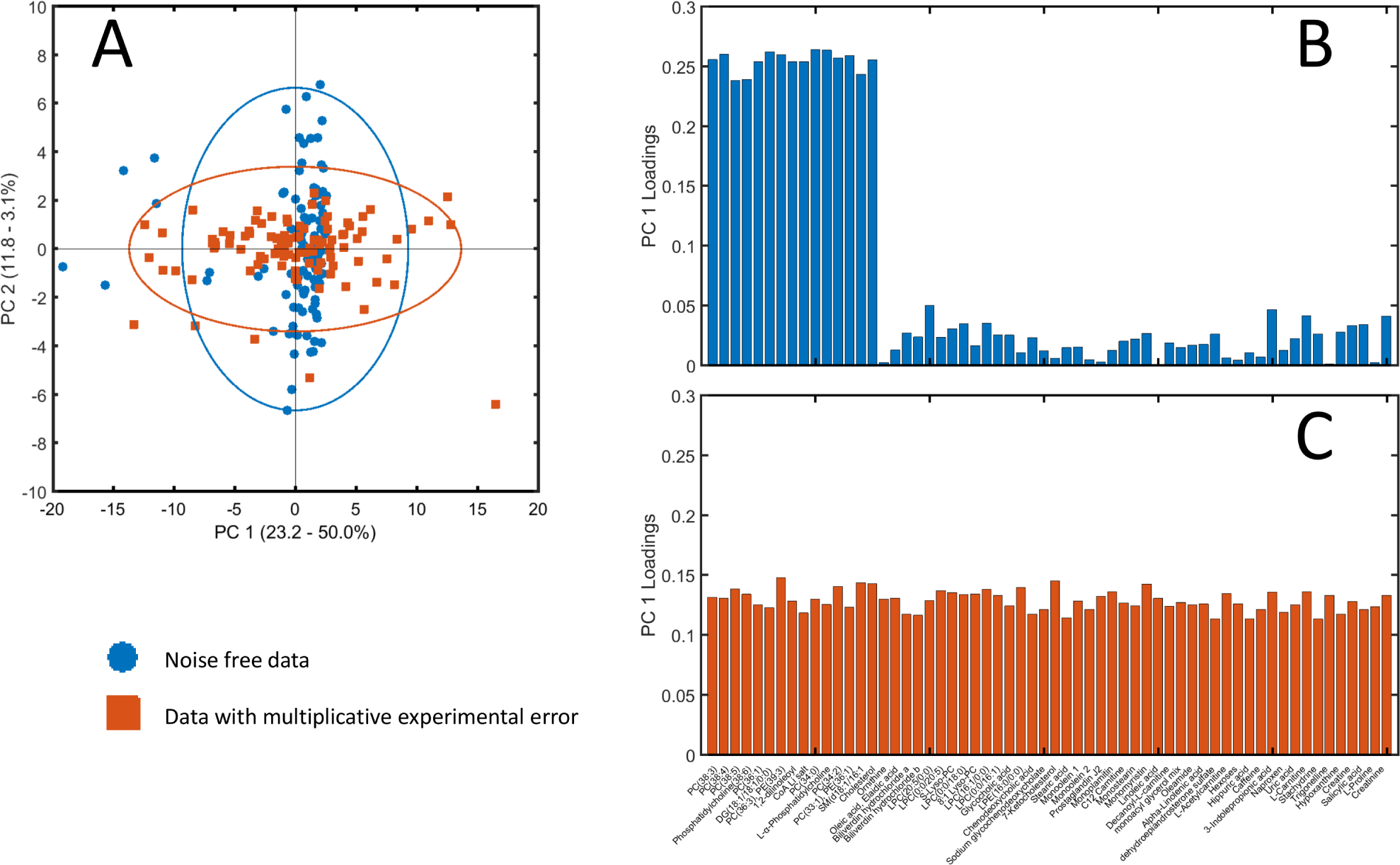
Consequences of multiplicative (correlated and uncorrelated) measurement error for data analysis. **A**: Scatter plot of the overlayed view of the first two components of two PCA models of simulated data sets; one without multiplicative error and one with multiplicative error. For visualization purposes, the scores are plotted in the same graph, but the subspaces spanned by the first two principal components for the two data sets are of course different. The labels on both axes also present the percentage explained variation for the two analyses. **B**: Loading plot for the error free data. **C**: Loading plot for the data with multiplicative error. See Material and Methods section 6.5.3 for details on the simulations.

### 2.3 Realistic error

The measurement process usually consists of different procedural steps and each step can be viewed as a different source of measurement error with its own characteristics, which sum to both additive and multiplicative error components as is the case of comprehensive *omics* measurements^27^. The model for this case is:

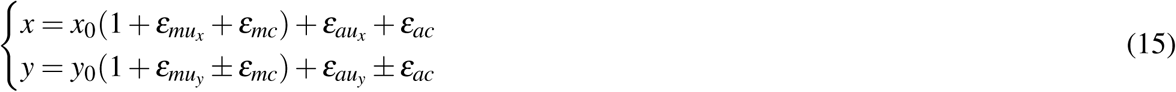

where all errors have been introduced before and are all assumed to be independent of each other and independent of the true (biological) signals (*x*_0_ and *y*_0_).

This realistic error model has a multiplicative as well as an additive component and also accommodates correlated and uncorrelated error. It is an extension of a much-used error model for analytical chemical data which only contains uncorrelated error^28^. From model (15) it follows that the error changes not only quantitatively but also qualitatively with changing signal intensity: the importance of the multiplicative component increases when the signal intensity increases, whereas the relative contribution of the additive error component increases when the signal decreases.

Since most of the measurements do not usually fall at the extremity of the dynamic range of the instruments used, the situation in which both additive and multiplicative error are important is realistic. For example, this is surely the case of comprehensive NMR and Mass Spectrometry measurements, where multiplicative errors are due to sample preparation and carry-over effect (in the case of MS) and the additive error is due to thermal error in the detectors^29^. To illustrate this we consider an NMR experiment where a different number of technical replicates are measured for five samples (Figure 4A and 4B). We are interested in establishing the correlation patterns across the (binned) resonances. For sake of simplicity we focus on two resonances, binned at 3.22 and 4.98 ppm. If one calculates the correlation using only one (randomly chosen) replicate per sample the resulting correlation can be anywhere between −1 and 1 (see Figure 4C.1). The variability reduces considerably if more replicates are taken and averaged before calculating the correlation (see Figure 4C), but there is still a rather large variation, induced by the limited sample size.

**Figure 4.**
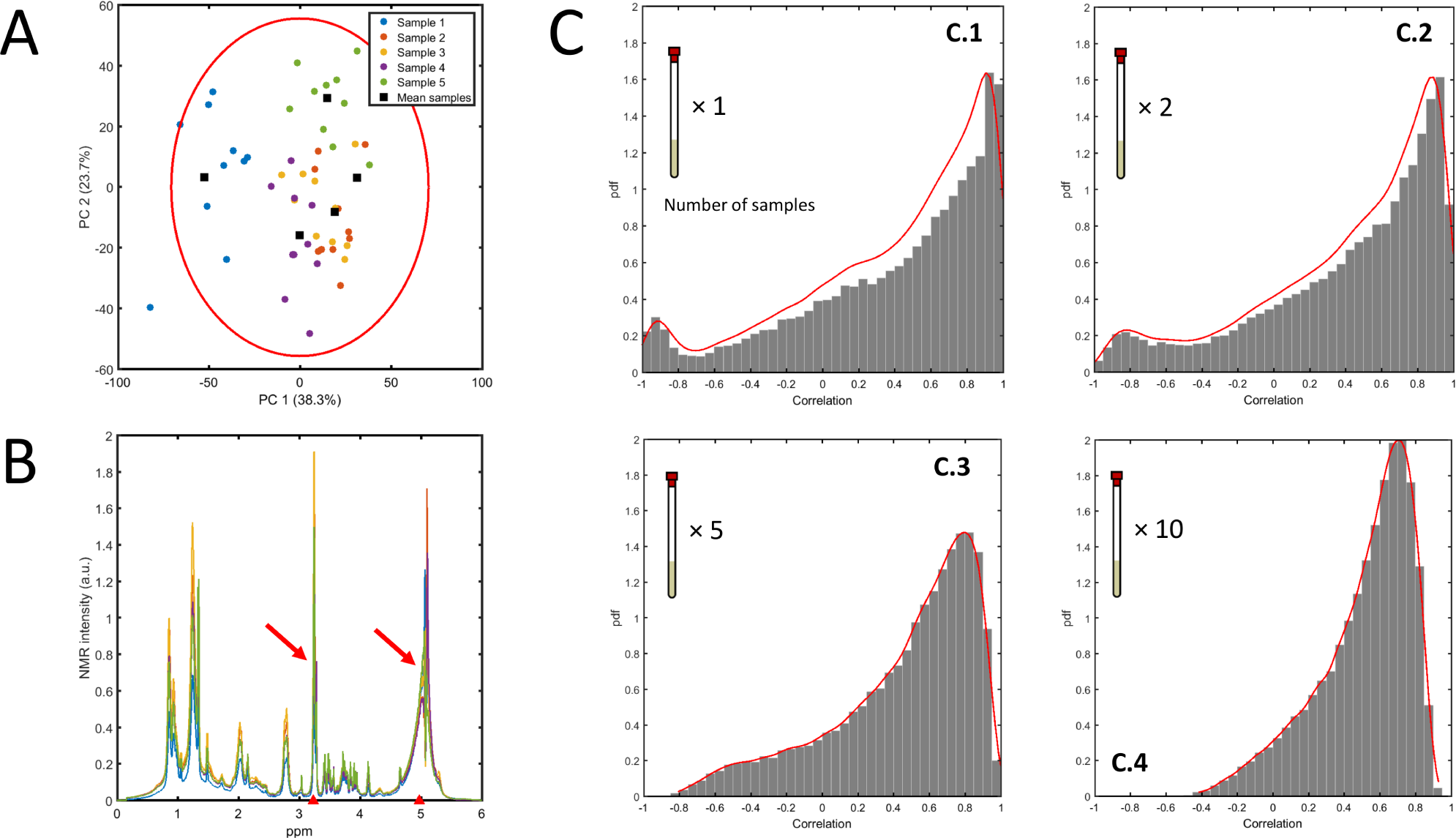
**A**: PCA plot of 5 different samples of fish extracts measured with technical replicates (10×) using NMR^29^. **B**: Overlap of the average binned NMR spectra of the 5 samples: the two resonances whose correlation is investigated are highlighted (3.23 and 4.98 ppm). **C**: Distribution of the correlation coefficient between the two resonances calculated, taking as input the average over different numbers of technical replicates (see inserts). See Material and Methods section 6.5.4 for more details on the estimation procedure.

Averaging across the technical replicates reduces variability among the sample means: however this not accompanied by an equal reduction in the variability of the correlation estimation. This is because the error is not taken into account in the calculation of the correlation coefficient.

## 3 Estimation of Pearson’s correlation coefficient in presence of measurement error

In the ideal case of an error free measurement where the only variability is due to intrinsic biological variation, ***ρ*** coincides with the true correlation ***ρ***_0_. If additive uncorrelated error is present, then ***ρ*** is given by Equations (9) and (10) which explicitly take into account the error component and it holds that ***ρ*** < ***ρ***_0_.

In the next Section we will derive analytical expressions, akin to Equations (9) and (10), for the correlation for variables sampled with measurement error (additive, multiplicative and realistic) as introduced in Section 2.

Before moving on, we define more specifically the error components. The error terms in models (11), and (15) are assumed to have the following distributional properties

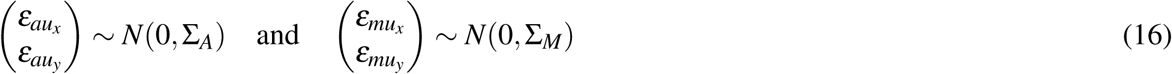

with

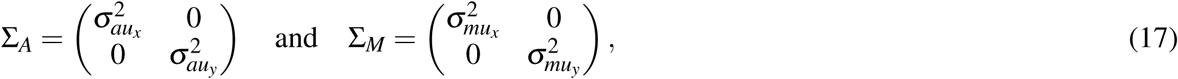

and

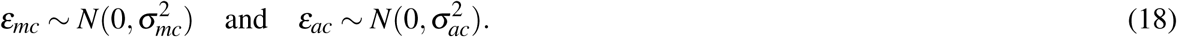

From definitions (16), (17) and (18) it follows that:

1. The expected value of the errors E[*ε*_*α*_] is zero:

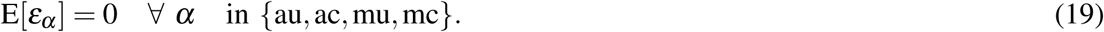
2. The covariance between *x*_0_ (*y*_0_) and the error terms is zero because *x*_0_ (*y*_0_) and errors are independent,

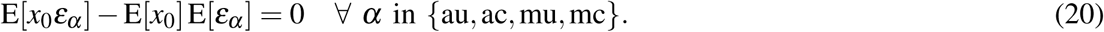
3. The covariance between the different error components is zero because the errors are independent from each other.

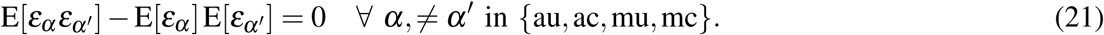

### 3.1 The Pearson correlation in the presence of additive measurement error

We show here a detailed derivation of the correlation among two variables *x* and *y* sampled under the additive error model (11). The variance for variable *x* (similar considerations hold for *y*) is given by

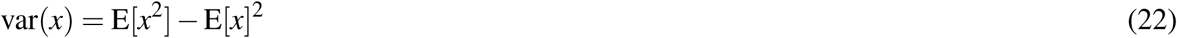

where

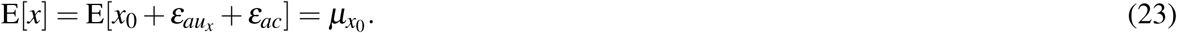

and

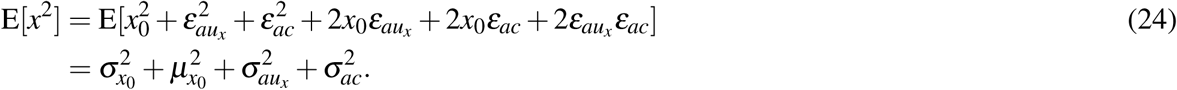

It follows that

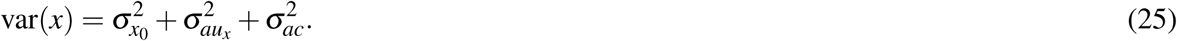

The covariance of *x* and *y* is

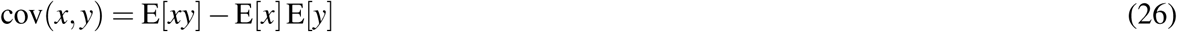

with

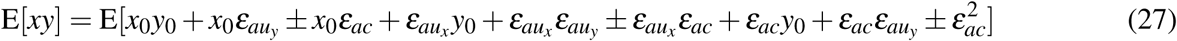

Considering (20) and (21), Equation (27) reduces to

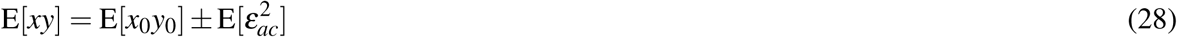

with

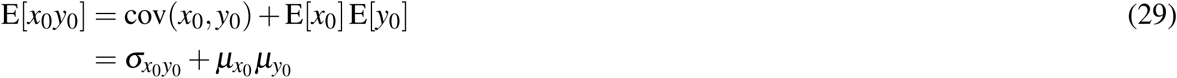

and

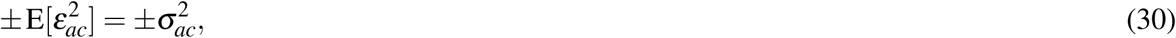

with ± depending on the sign of the measurement error correlation. From Equations (23), (28), (29), and (30) it follows

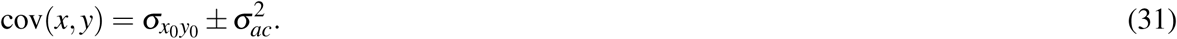

Plugging (25) and (31) into (6) and defining the attenuation coefficient *A*^*a*^

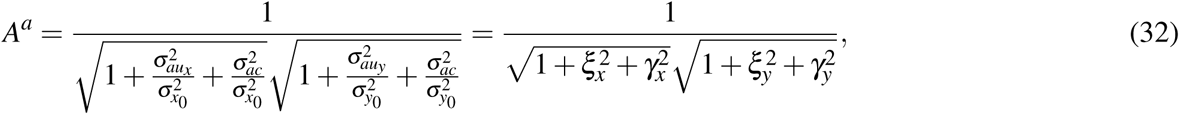

where 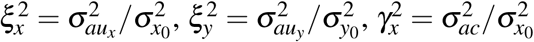 and 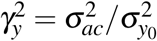; the superscript *a* in *A*^*a*^ stands for *a*dditive.

The Pearson correlation in presence of additive measurement error is obtained as:

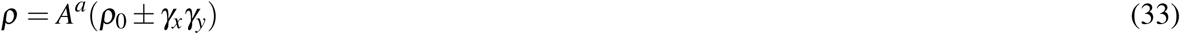

where the sign ± signifies positively and negatively correlated error.

The attenuation coefficient *A*^*a*^ is a decreasing function of the measurement error ratios, that is, the ratio between the variance of the uncorrelated and the correlated error to the variance of the true signal. Compared to Equation (9), in formula (33) there is an extra additive term related to the correlated measurement error expressing the impact of the correlated measurement error relative to the original variation. In the presence of only uncorrelated error (*i.e.* 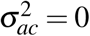), Equation (33) reduces to the Spearman’s formula for the correlation attenuation given by (9) and (10). As previously discussed, in this case the correlation coefficient is always biased towards zero (attenuated).

Given the true correlation ***ρ***_0_, the expected correlation coefficient (33) is completely determined by the measurement error ratios. Assuming the errors on *x* and *y* to be the same (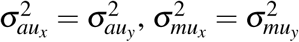, an assumption not unrealistic if *x* and *y* are measured with the same instrument and under the same experimental conditions during an *omics* comprehensive experiment) and taking for simplicity 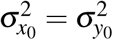, then *ξ*_*x*_ = *ξ*_*y*_ = *ξ* and *γ*_*x*_ = *γ*_*y*_ = *γ* and Equation (33) can be simplified to:

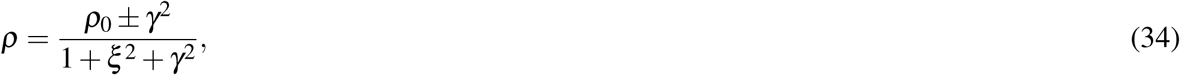

and ***ρ*** can be visualized graphically as a function of the uncorrelated and correlated measurement error ratios *ξ* and *γ* as shown in Figure 5.

**Figure 5.**
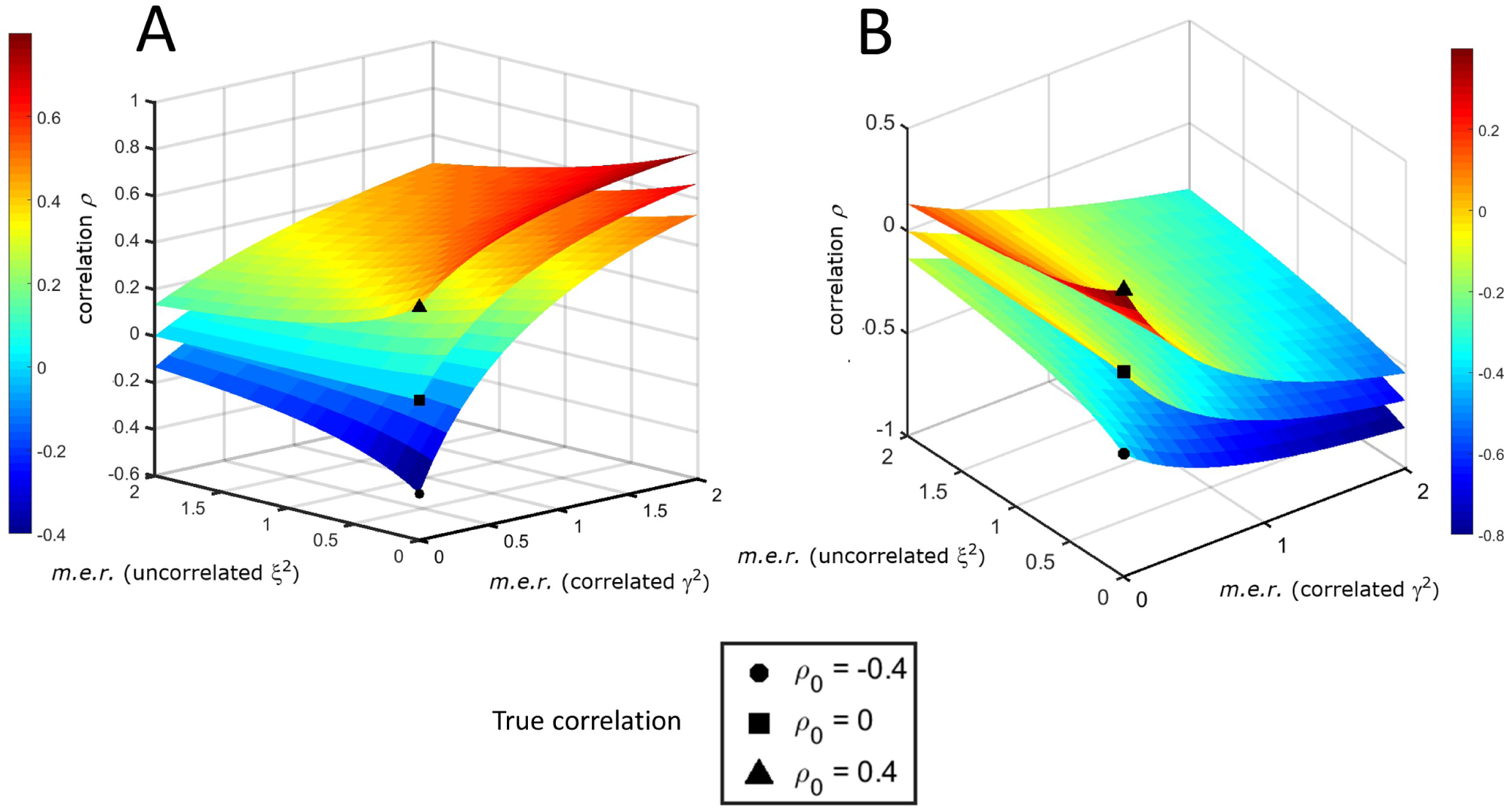
The expected correlation coefficient ***ρ*** in the presence of additive measurement error as a function of the uncorrelated (*ξ* ^2^) and correlated (*γ*^2^) measurement error ratios (*m.e.r.*) for different values of the true correlation ***ρ***_0_. **A**: Positively correlated error. **B**: Negatively correlated error.

In the presence of positively correlated error, the correlation ***ρ*** is attenuated towards 0 if the uncorrelated error increases and inflated if the additive correlated error increases (Figure 5A, which refers to Equation (34)) when ***ρ***_0_ > 0. If ***ρ***_0_ < 0 the distortion introduced by the correlated error can be so severe that the correlation ***ρ*** can become positive. When the error is negatively correlated (Figure 5B), the correlation ***ρ*** is biased towards 0 when ***ρ***_0_ > 0 (and can change sign), while it can be attenuated or inflated if ***ρ***_0_ < 0.

A set of rules can be derived to describe quantitatively the bias of ***ρ***. For positively correlated measurement error (for negatively correlated measurement error see Section 6. 6.2) if the true correlation ***ρ***_0_ is positive the correlation ***ρ*** is always strictly positive: this is shown on Figure 6A where the relationship between ***ρ*** and ***ρ***_0_ is shown by means of Monte Carlo simulation (see Figure caption for more details). The magnitude of ***ρ*** (‖***ρ***‖) depends on how *A*^*a*^ (for readability in the following equations we will use *A*) and the additive term *γ*_*x*_*γ*_*y*_ > 0 compensate each other. In particular when ***ρ***_0_ > 0

**Figure 6.**
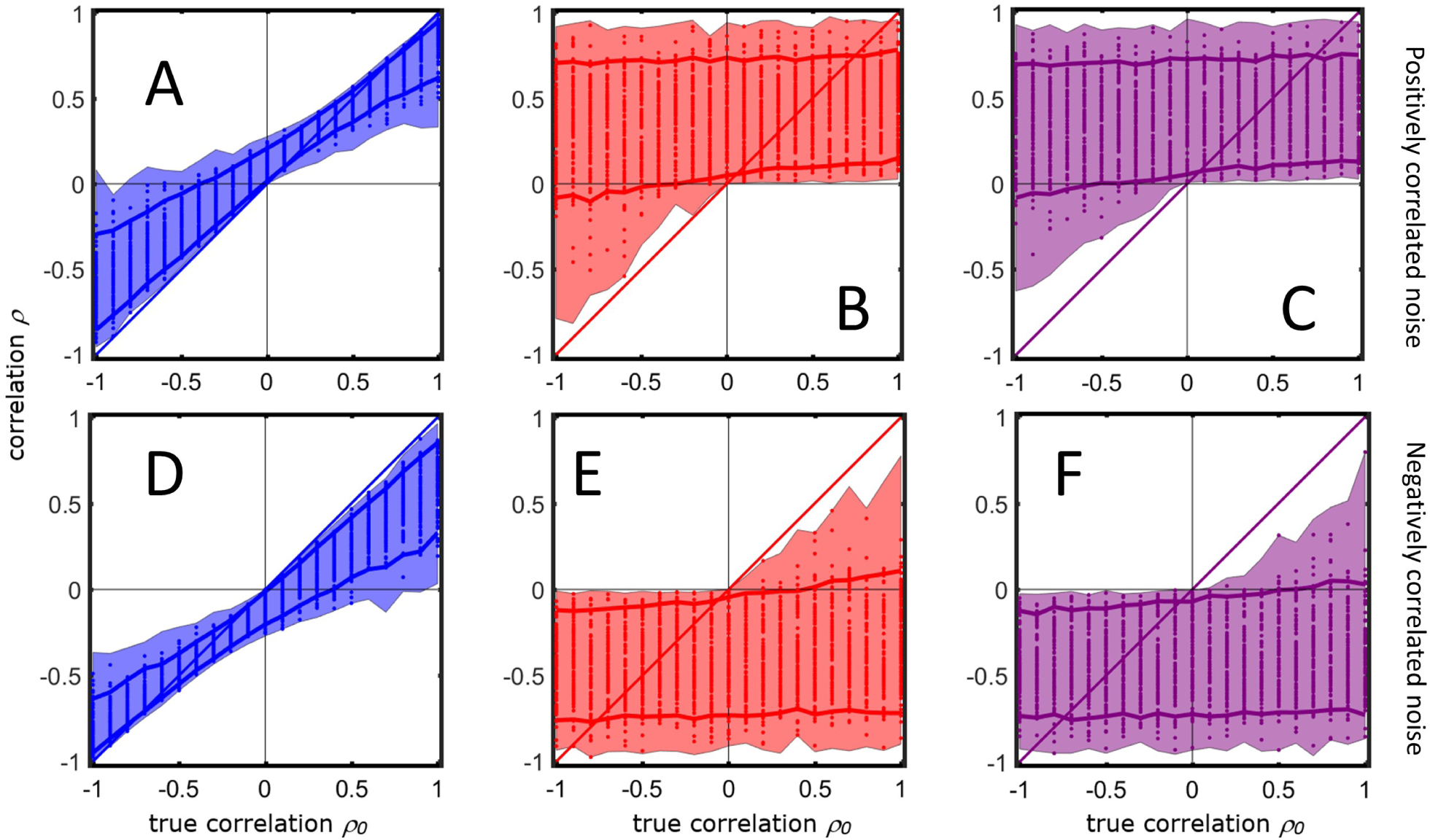
Calculations of the correlation coefficient ***ρ*** (40) as a function of the different realizations of the signal means and the size of the error components for different values of the true correlation ***ρ***_0_. The shadowed area encloses the maximum and the minimum of the values of ***ρ*** calculated in the simulation using the different error models. The dots represent the realized values of ***ρ*** (only 100 of 10^5^ Monte Carlo realizations for different values of the variances of error component are shown). The solid lines represent the 5-th and the 95-th percentiles of the observed values. **A**: Additive measurement error with positive correlated error. **B**: Multiplicative measurement error with positive correlated error. **C** Realistic case with both additive and multiplicative measurement error with positive correlated error. **D**: Additive measurement error with negative correlated error. **E**: Multiplicative measurement error with negative correlated error. **F**: Realistic case with both additive and multiplicative measurement error with negative correlated error. For more details on the simulations see Material and Methods section 6.5.5.

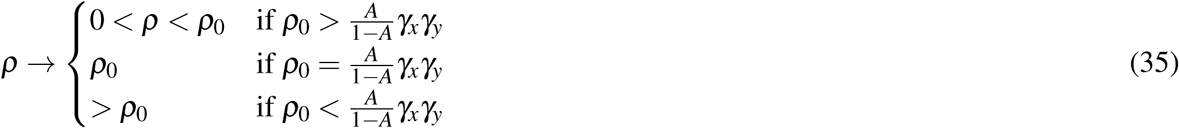

This means that ***ρ*** is always a biased estimator of the true correlation ***ρ***_0_, with the exception of the second case which happens only for specific values of *γ* and ***ρ***_0_. This is unlikely to happen in practice.

If ***ρ***_0_ < 0 it holds that

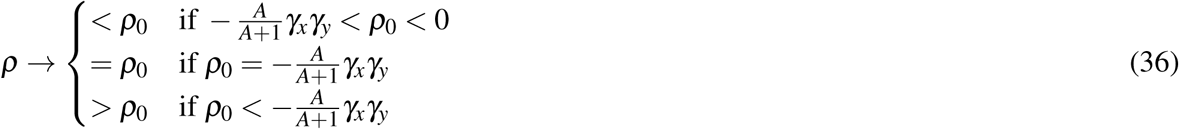

The interpretation of Equation (36) is similar to that of Equation (35) but additionally, the correlation coefficient can even change sign. In particular, this happens when

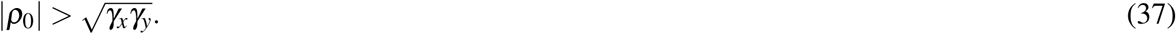

The terms 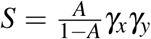 and 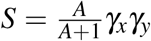 in Equations (35), (36), (71) and (72) describe limiting surfaces *S* of ***ρ***_0_ values delineating the regions of attenuation and inflation of the correlation coefficient *ρ.* As can be seen from Figure 7, these surfaces are not symmetric with respect to zero correlation, indicating that the behavior of ***ρ*** is not symmetric around 0 with respect to the sign of ***ρ***_0_ and of the correlated error.

**Figure 7.**
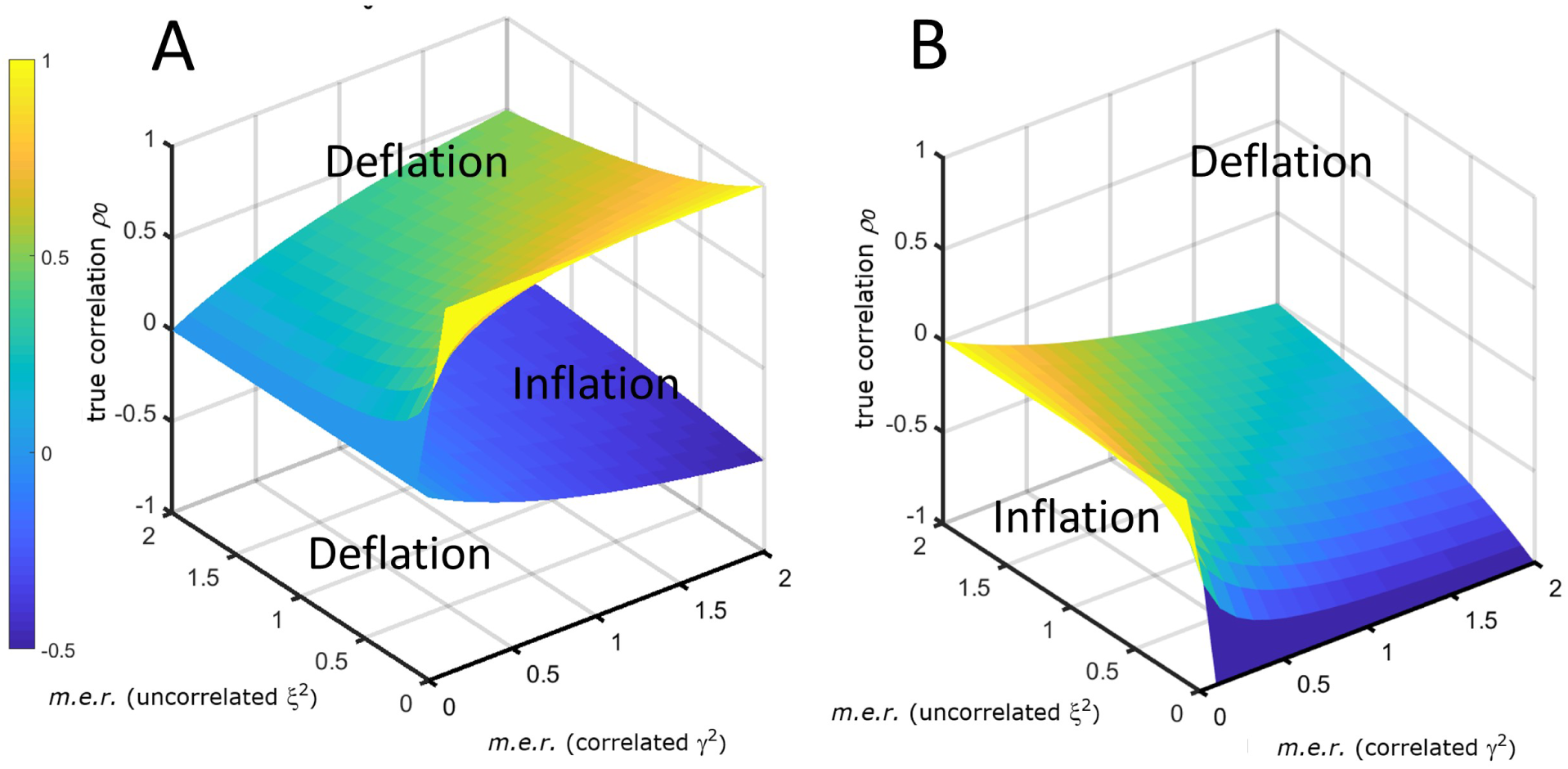
Limiting surfaces *S* for the inflation and deflation region of the correlation coefficient in presence of additive measurement error. The surfaces are a function of the uncorrelated (*ξ* ^2^) and correlated (*γ*^2^) measurement error ratios (*m.e.r.*). **A**: *S* in the case of positively correlated error. **B**: *S* for negatively correlated error. The plot refers to ***ρ*** defined by Equation (34) with 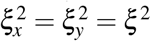 and 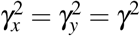.

### 3.2 The Pearson correlation in presence of multiplicative measurement error

The correlation in the presence of multiplicative error can be derived using similar arguments and detailed calculations can be found in Section 6.1.1. Here we only state the main result:

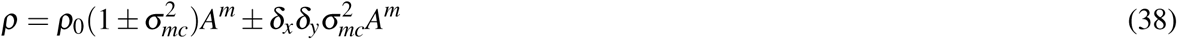

with 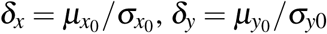 (biological signal to biological variation ratios) and *A*^*m*^ is the attenuation coefficient (the superscript *m* stands for *m*ultiplicative):

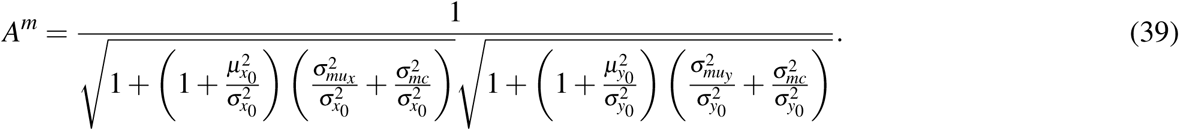

In this case, the correlation coefficient depends explicitly on the mean of the variables, as an effect of the multiplicative nature of the error component.

Our simulations show that if the signal intensity is not too large, the correlation can change sign (as shown in Figure 6B); if the signal intensity is very large the multiplicative error will have a very large effect and if the correlated error is positive the expected correlation ***ρ*** will also be positive, and will be negative if the error are negatively correlated. but simulations cannot be exhaustive (as shown in Figure 6B.)

### 3.3 The Pearson correlation in presence of realistic measurement error

When both additive and multiplicative error are present, the correlation coefficient is a combination of formula (33) and (38) (see Section 6.1.2 for detailed derivation):

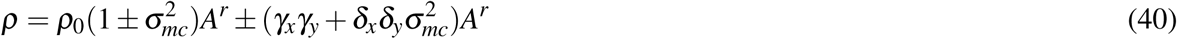

Where the *γ* and *δ* parameters have been previously defined for the additive and multiplicative case. *A*^*r*^ is the attenuation coefficient (the superscript *r* stands for *r*ealistic):

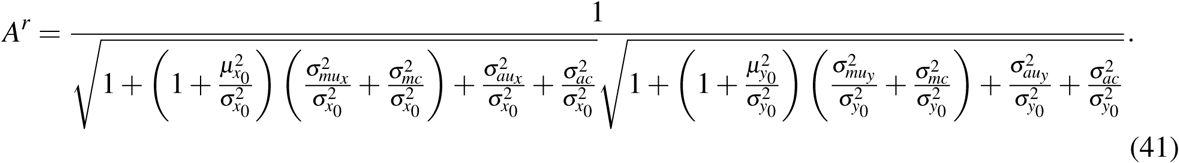

General rules governing the sign of the numerator and denominator in Equation (40) cannot be determined since it depends on the interplay of the six error components, the true mean and product thereof. Within the parameter setting of our simulations, the results presented in Figures 6C show that the behavior of ***ρ*** under error model 15 is qualitatively similar to that in presence of only multiplicative error. However different behavior could be emerge with different parameter settings.

### 3.4 Generalized correlated error model

The error models presented in Equations (11), (13), and (15) assume a perfect correlation of the correlated errors, since the correlated error terms *ε*_*ac*_ appear simultaneously in both *x* and *y*; the same hold true for *ε*_*mc*_. A more general model that accounts for different degrees of correlation between the error components can be obtained by modifying the model (15) (other cases are treated in Section 6.3.) to

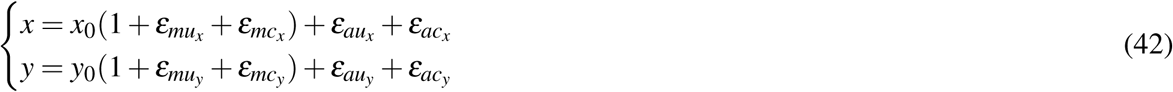

where the correlated error components 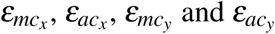 and 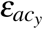 are distributed as

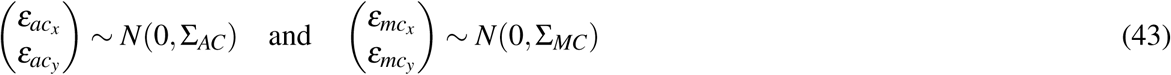

with

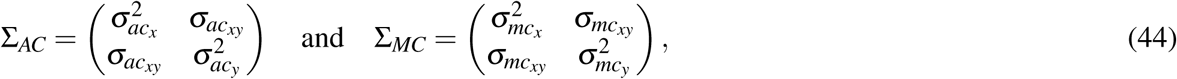

where 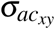 is the covariance between error term 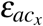 and 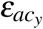 and 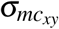 is the covariance between error term 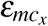 and 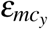.

It is possible to derive expression for the correlation coefficient under the model (43) as shown in Section 3.1 and in the Section 6.1.1 and 6.1.2. The only difference is that under this model the terms 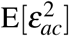 and 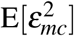 in Equations (27), (58), (65), and (66) are replaced by 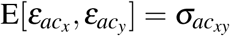 and 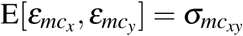, respectively.

From the definition of covariance it follows that

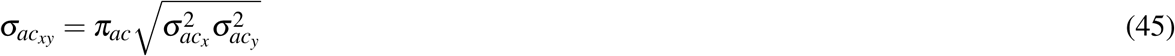

and

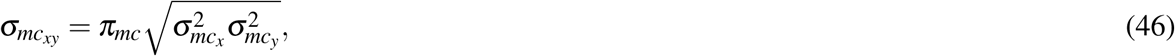

where *π*_*ac*_ and *π*_*mc*_ are the correlations among the error terms for which it holds −1 ≤ *π*_*mc*_ ≤ 1 and −1 ≤ *π*_*mc*_ ≤ 1. If *π*_*ac*_ and *π*_*mc*_ are negative the errors are negatively correlated. Equation (40) becomes now:

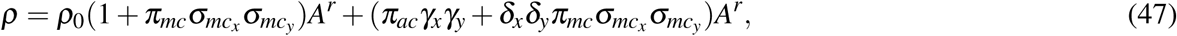

with 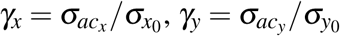, and

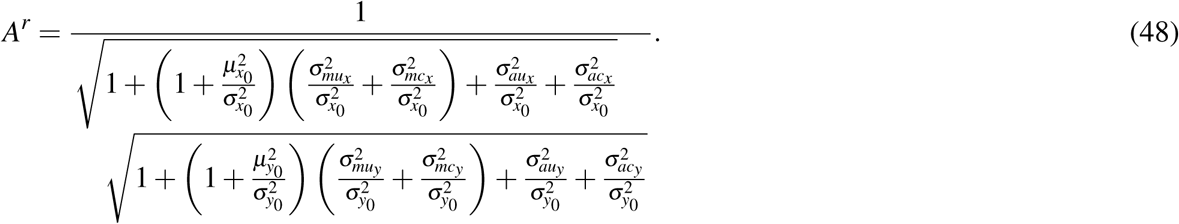

This model generalizes the correlation coefficient among *x* and *y* from Equation (40) to account for different strength of the correlation among the correlated error components. All considerations discussed in the previous sections do apply also to this model. Expressions for ***ρ*** in the case of additive and multiplicative error can be found in the Section 6.3.1 and 6.3.2.

By setting 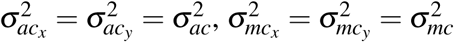, and *π*_*ac*_ = *π*_*mc*_ = 1 (perfect correlation), model (40) is obtained, and similarly models (33) and (38).

## 4 Correction for correlation bias

Because virtually all kinds of measurement are affected by measurement error, the correlation calculated from sampled data is distorted to some degree depending on the level of the measurement error and on its nature. We have seen that experimental error can inflate or deflate the correlation and that ***ρ*** (and hence its sample realization *r*) is almost always a biased estimation of the true correlation ***ρ***_0_. An estimator that gives a theoretically unbiased estimate of the correlation coefficient between two variables *x* and *y* taking into account the measurement error model can be derived. For simple uncorrelated additive error this is given by the Spearman’s formula (49): this is a known results which in the past has been presented and discussed in many different fields^16–19^. To obtain similar correction formulas for the error models considered here it is sufficient to solve for ***ρ***_0_ from the defining Equations (33), (38) and (40). The correction formulas are as follows (the ± indicates positive and negatively correlated error):

1. Correction for simple additive error (only uncorrelated error):

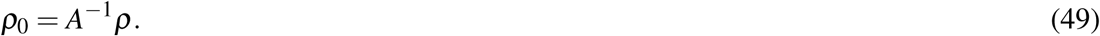
2. Correction for additive error:

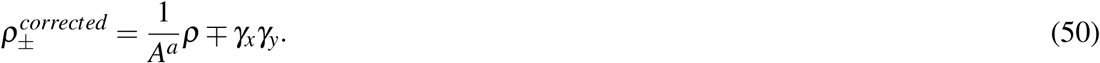
3. Correction for multiplicative error:

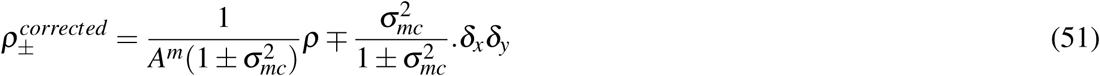
4. Correction for realistic error:

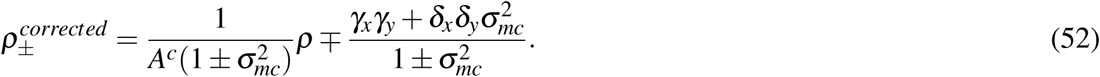

In practice, to obtain a corrected estimation of the correlation coefficient ***ρ***_0_, the ***ρ*** is substituted by *r* in (50), (51) and (52), which is the sample correlation calculated from the data. The effect of the correction is shown, for the realistic error model (15), in Figure 8 where the true know error variance components have been used. It should be noted that it is possible that the corrected correlation exceeds ±1.0. This phenomenon has already been observed and discussed^16, 30^: it is due to the fact that the sampling error of a correlation coefficient corrected for distortion is greater than would be that of an uncorrected coefficient of the same size (at least for the uncorrelated additive error^4, 18, 31^). When this happens the corrected correlation can be rounded to ±1.0^19, 31^.

**Figure 8.**
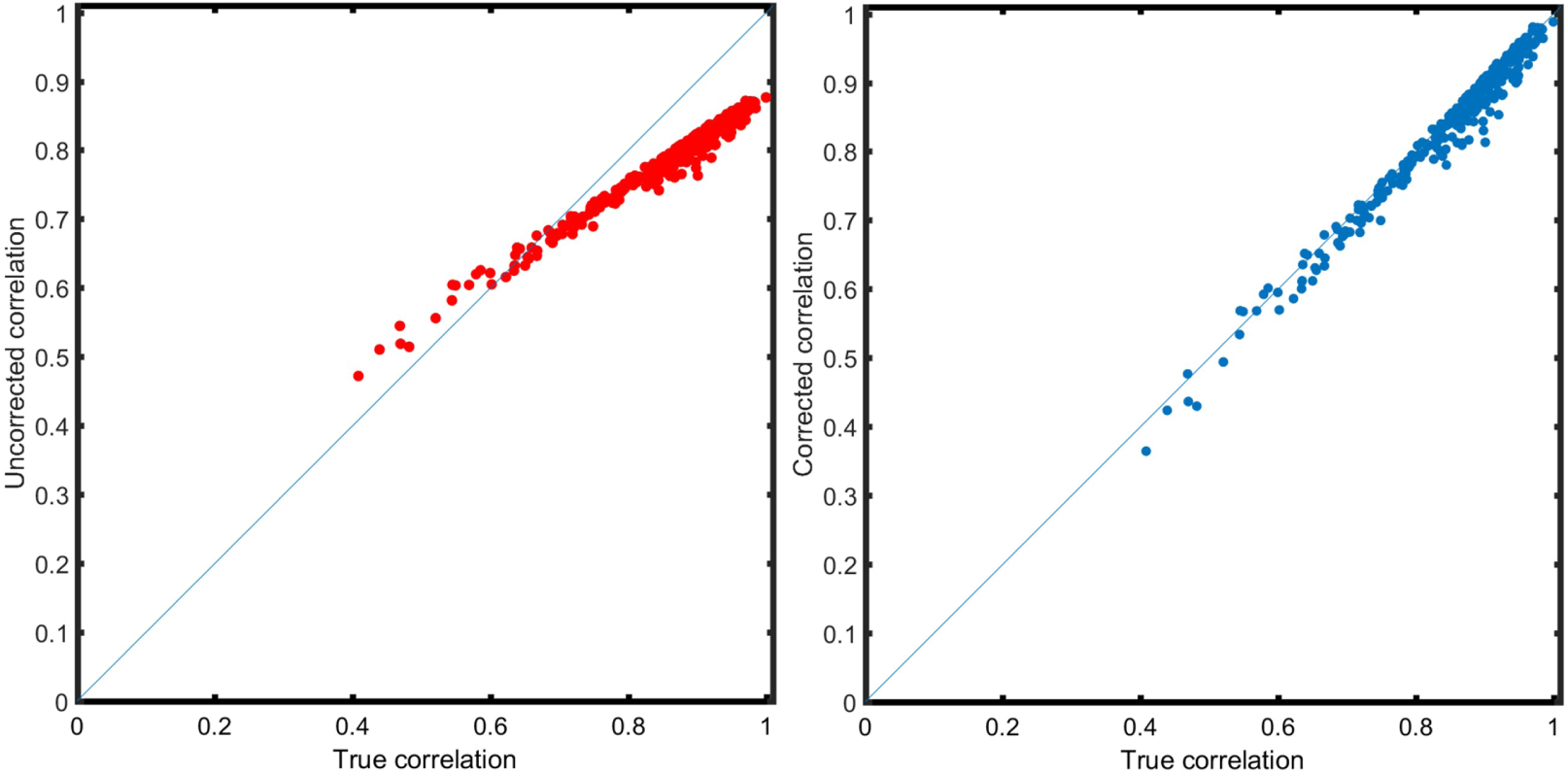
Correction of the distortion induced by the realistic measurement error (see Equation (15)). **A**: Pairwise correlations ***ρ*** among 25 metabolites calculated from simulated data with additive and multiplicative measurement error *vs* the true correlation ***ρ***_0_. **B**: Corrected correlation coefficients using Equation 52 and using the known error variance components. See Section 6.5.6 for details on the data simulation.

### 4.1 Estimation of the error variance components

Simulations shown in Figure 8 have been performed using the known parameters for the error components used to generate the data. In practical applications the error components needs to be estimated from the measured data and the quality of the correction will depend on the accuracy of the error variance estimate.

The case of purely additive uncorrelated measurement error 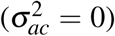 has been addressed in the past^18, 19, 32^: in this case the variance components 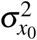 and 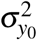 can be substituted with their sample estimates (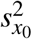 and 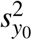) obtained from measured data, while the error variance components (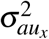 and 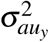) can be estimated if an appropriate experimental design is implemented, *i.e.* if *n* replicates are measured for each observation.

Unfortunately, there is no simple and immediate approach to estimate the error component in the other cases when many variance components need to be estimated (6 error variances in the case of error model (15) and 8 in the case of the generalized model (42), to which the estimations of *π*_*mc*_ and *π*_*ac*_) must be added).

Different approaches can be foreseen to estimate the error components which is not a trivial task, including the use of (generalized) linear mixed model^33, 34^, error covariance matrix formulation^29, 35, 36^ or common factor analysis factorization^37^. None of these approaches is straightforward and require some extensive mathematical manipulations to be implemented; an accurate investigation of the simulation of the error component is outside the scope of this paper and will presented in a future publication.

## 5 Discussion

Since measurement error cannot be avoided, correlation coefficients calculated from experimental data are distorted to a degree which is not known and that has been neglected in life sciences applications but can expected to be considerable when comprehensive omics measurement are taken.

As previously discussed, the attenuation of the correlation coefficient in the presence of additive (uncorrelated) error has been known for more than one century. The analytical description of the distortion of the correlation coefficient in presence of more complex measurement error structures (Equations (33), (38) and (40)) has been presented here for the first time to the best of our knowledge.

The inflation or attenuation of the correlation coefficient depends on the relationship between the value of true correlation ***ρ***_0_ and the error component. In most cases in practice, ***ρ*** is a biased estimator for ***ρ***_0_. In absence of correlated error, there is always attenuation; in the presence of correlated error there can also be increase (in absolute value) of the correlation coefficient. This has also been observed in regression analysis applied to nutritional epidemiology and it has been suggested that correlated error can, in principle, be used to compensate for the attenuation^38^. Moreover, the distortion of the correlation coefficient has also implications for hypothesis testing to assess the significance of the measured correlation *r*.

To illustrate the counterintuitive consequences of correlated measurement error consider the following. Suppose that the true correlation is null. In that case, Equations (33), (38) and (40) reduce to

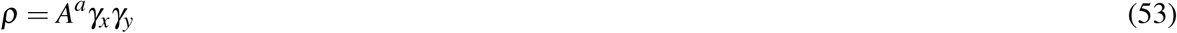

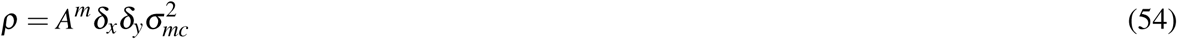

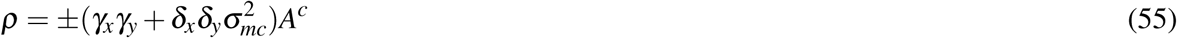

which implies that the correlation coefficient is not zero. Moreover, in real-life situations there is also sampling variability superimposed on this which may in the end result in estimated correlations of the size as found in several omics applications (in metabolomics observed correlations are usually lower than 0.6^10, 22^; similar patterns are also observed in transcriptomics^39, 40^) while the true biological correlation is zero.

The correction equations presented need the input of estimated variances. Such estimates also carry uncertainty and the quality of these estimates will influence the quality of the corrections. This will be the topic of a follow-up paper. Prior information regarding the sizes of the variance components would be valuable and this points to new requirements for system suitability tests of comprehensive measurements. In metabolomics, for example, it would be worthwhile to characterize an analytical measurement platform in terms of such error variances including sizes of correlated error using advanced (and to be developed) measurement protocols.

Distortion of the correlation coefficient has implications also for experimental planning. In the case of additive uncorrelated error, the correction depends explicitly on the sample size *N* used to calculate *r* and on the number of replicates *n*_*x*_, *n*_*y*_ used to estimate the intraclass correlation (*i.e.* the error variance components): since in real life the total sample size *N* × (*n*_*x*_ + *n*_*y*_) is fixed, there is a trade off between the sample size and the number of replicates that can be measured and the experimenter has to decide whether to increase *N* or *n*_*x*_.

The results presented here are derived under the assumption of normality of both measurement and measurement errors. If *x*_0_ and *y*_0_ are normally distributed, then *x* and *y* will be, in presence of additive measurement error, normally distributed, with variance given by (12). For multiplicative and realistic error the distribution of *x* and *y* will be far from normality since it involves the distribution of the product of normally distributed quantities which is usually not normal^41^. It is known that departure from normality can result in the inflation of the correlation coefficient^42^ and in distortion^43^ of its (sampling) distribution and this will add to the corruption induced by the measurement error.

We think that in general correlation coefficients are trusted too much on face value and we hope to have triggered some doubts and pointed to precautions in this paper.

## 6 Material and Methods

### 6.1 Mathematical calculations

#### 6.1.1 Derivation of ρ in presence of multiplicative measurement error

In presence of purely multiplicative error it holds

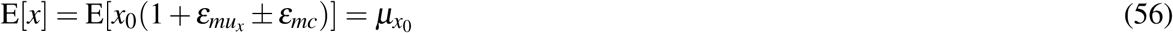

and

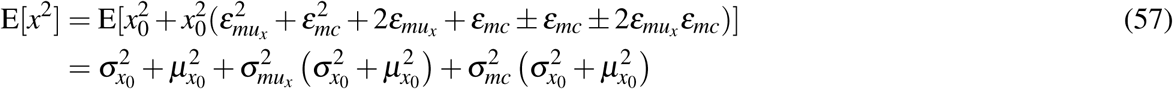

using (19) – (21) main text to calculate the expectation of the cross terms. For E[*xy*] it holds

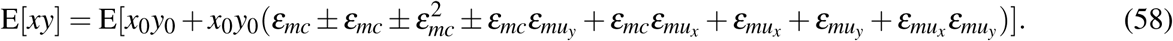

Because of the independence of *x*_0_, *y*_0_ and the error terms, the expectations of all cross terms is null except

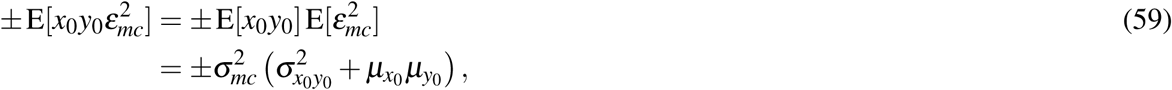

where E[*x*_0_*y*_0_] is given by Equation (29). Plugging (56), (57) and (58) in (6), the expected correlation coefficient is

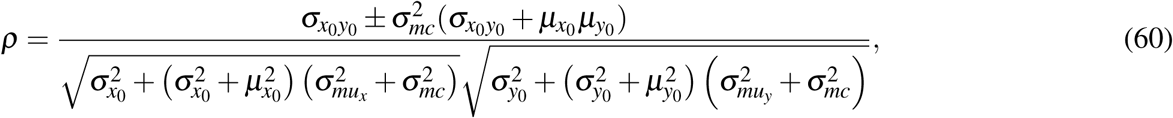

and it can re-written as (38) by setting 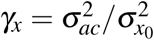 and 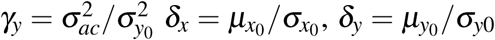 and defining the attenuation coefficient *A*^*m*^ (39).

#### 6.1.2 Derivation of ρ in presence of realistic measurement error

To simplify calculations we set

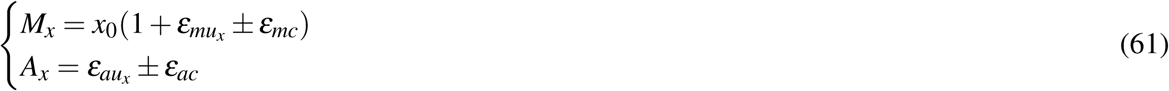

and similarly we define *M*_*y*_ and *A*_*y*_ for variable *y*. It holds

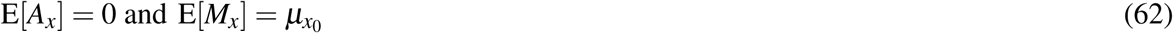

and

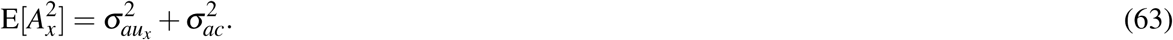

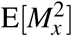 is given by Equation (57). Because error components are independent and with zero expectation (see Equations (19)-(21)) it holds

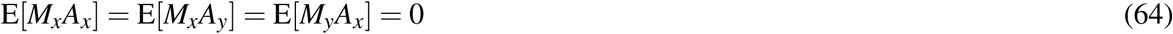

and

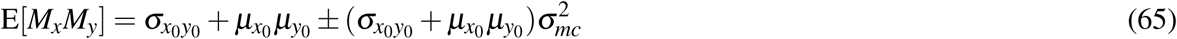

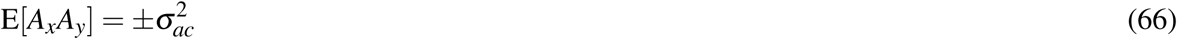

It follows that

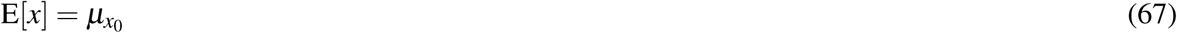

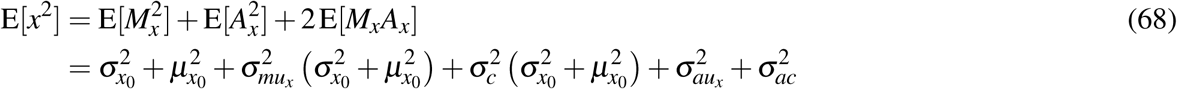

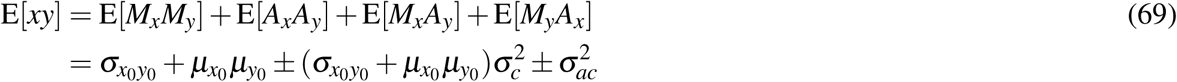

Plugging (67), (68), and (69) into (6) one gets the expression for the correlation coefficient in presence of additive and multiplicative measurement error:

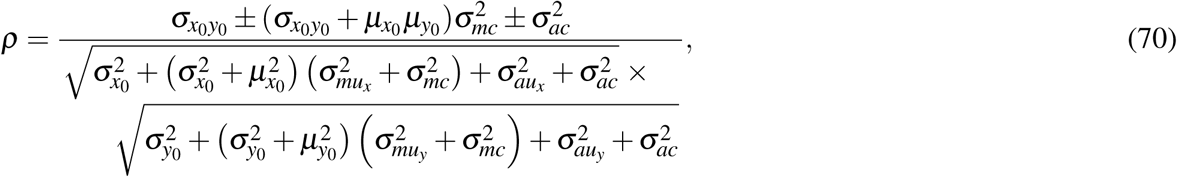

that can re-written as (40) by using previously defined *γ*_*x*_, *γ*_*y*_, *δ*_*x*_ and *δ*_*y*_ and defining the attenuation coefficient *A*^*c*^ (41).

### 6.2 Behavior of ***ρ*** in the case of additive negatively correlated error

For negative correlated error, when the true correlation is positive

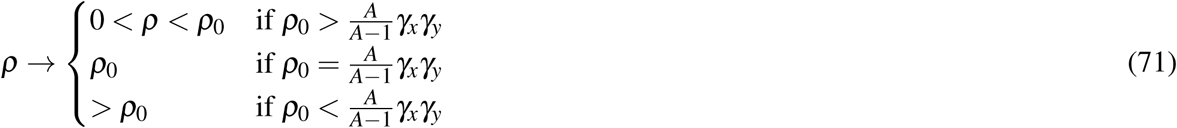

Since 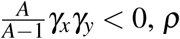, is always smaller than the true correlation. When the true correlation is negative (***ρ***_0_ the expected correlation is always negative, but it can be, in absolute value, smaller or larger than the absolute value of the true correlation:

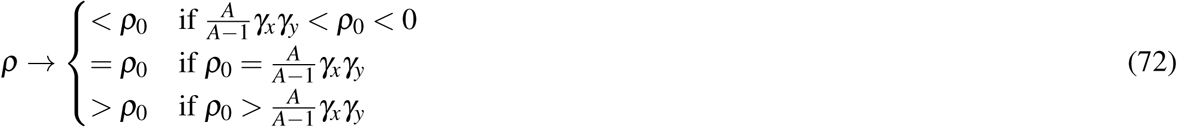

### 6.3 Correlation coefficient under the generalized error model

#### 6.3.1 Additive error

Under the generalized additive correlated error model

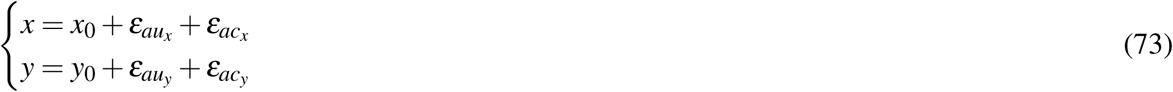

with 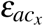 and 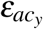 defined in Equations (43), the correlation coefficient can be expressed as:

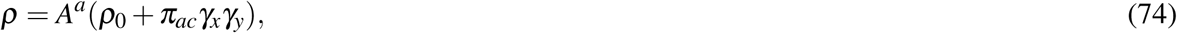

with 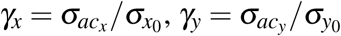, and

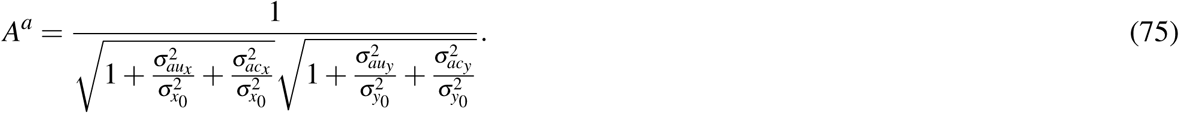

#### 6.3.2 Multiplicative error

Under the generalized multiplicative error model

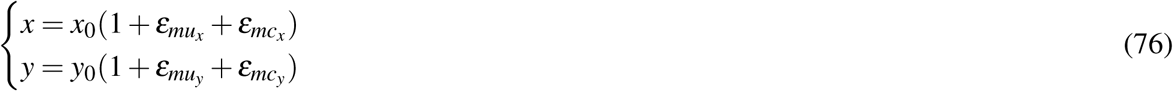

with 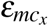 and 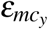 defined in Equations (43), the correlation coefficient can be expressed as:

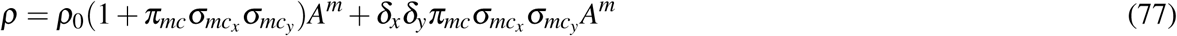

with

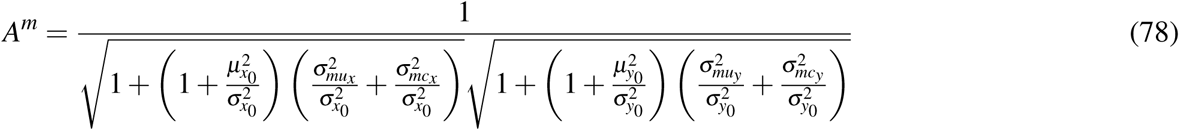

#### 6.3.3 General realistic error

Formulas for the correlation coefficient under the generalized realistic correlated error model are to be found in the main text in Equations (47) and (48).

### 6.4 Correction of the correlation coefficient under the generalized correlated error model

#### 6.4.1 Additive error

Under the generalized additive correlated error model the corrected correlation coefficient is

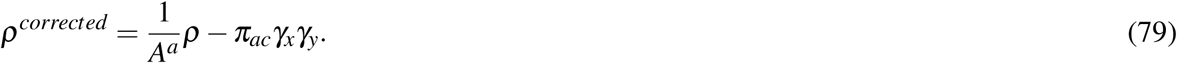

#### 6.4.2 Multiplicative error

Under the generalized multiplicative correlated error model the corrected correlation coefficient is

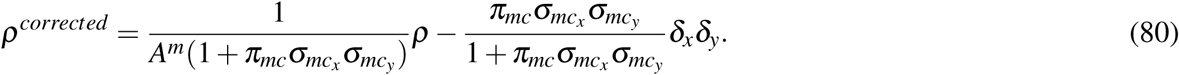

#### 6.4.3 Realistic error

Under the generalized realistic correlated error model the corrected correlation coefficient is

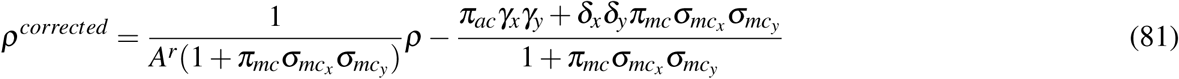

### 6.5 Simulations

We provide here details on the simulation performed and shown in Figures 1 – 4, 6 and 8.

#### 6.5.1 Simulations in Figure 1

*N* = 100 realizations of two variables *x* and *y* were generated under model with additive uncorrelated measurement error (11), with 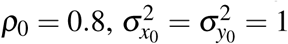 and *µ* = (100, 100). Error variance components were set to 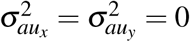 and to 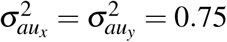 (Panel A).

#### 6.5.2 Simulations in Figure 2

The time concentrations profiles *P*_1_(*t*), *P*_2_(*t*) and *P*_3_(*t*) of three hypothetical metabolites P1, P2 and P3 are simulated using the following dynamic model

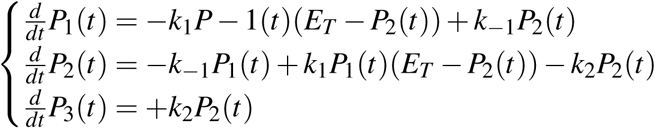

which is the model of an irreversible enzyme-catalyzed reaction described by Michaelis-Menten kinetics. Using this model, *N* = 100 concentration time profiles for P1, P2 and P3 were generated by solving the system of differential equations after varying the kinetic parameters *k*_1_, *k*_−1_ and *k*_2_ by sampling them from a uniform distribution. For the realization of the *j*th concentration profile

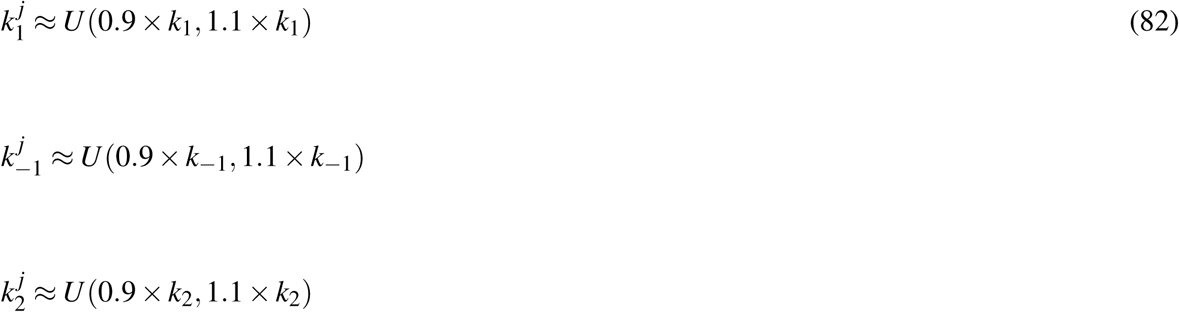

with population values *k*_1_ = 30, *k*_−1_ = 20, *k*_2_ = 10, and *E*_*T*_ = 1. Initial conditions were set to 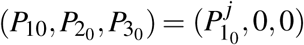 with 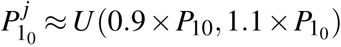 and 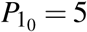. All quantities are in arbitrary units. Time profiles were sampled at *t* = 0.4 *a.u* and collected in a data matrix **X**_0_ of size 100 × 3. The variability in data matrix **X**_0_ is given by biological variation. The concentration time profiles of P1, P2 and P3 shown in Panel A are obtained using the population values for the kinetic parameters and for the initial conditions.

Additive uncorrelated and correlated measurement error is added on **X**_0_ following model (11) where P1, P2 and P3 in **X**_0_ play the role of *x*_0_, *y*_0_ and of an additional third variable *z*_0_ which follows a similar model. The variance of the error component was varied in 50 steps between 0 and 25% of the sample variance 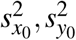 and 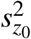 calculated from **X**_0_. The variance of the correlated error was set to 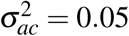 in all simulations. Pairwise Pearson correlations *r*_*i, j*_ with *i, j* = {*P*1, *P*2, *P*3} were calculated for the error free case **X**_0_ and for data with measurement error added. 100 error realization were simulated for each error value and the average correlation across the 100 realization is calculated and it is shown in Panel B.

The “mini” metabolite-metabolite association networks shown in Panel C are defined by first taking the Pearson correlation *r*_*i j*_ among P1, P2 and P3 and then imposing a threshold on *r* to define the Connectivity matrix *A*_*i j*_

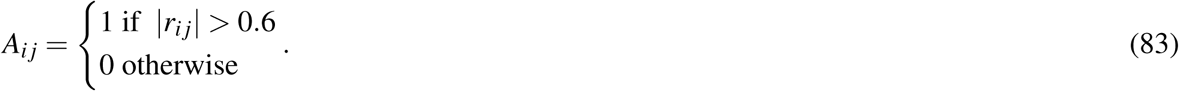

For more details see reference^10^.

#### 6.5.3 Simulations in Figure 3

Principal component analysis was performed on a 100 × 133 experimental metabolomic data set (see Section 6.6 for a description). The 15 variables with the highest loading (in absolute value) and the 45 variables with the smallest loading (in absolute value) on the first principal component where selected to form a 100 × 60 data set **X**_0_ (we call this now the error free data, as if it only contained biological variation). On this subset a new a principal component analysis was performed. Then multiplicative correlated and uncorrelated measurement error was added on **X**_0_. The variance of the additive error was set 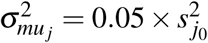 with *j* = 1, 2, …, 60 where 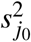 is the variance calculated for the *j*th column of **X**_0_, *i.e.*, the biological variance. The variance of the correlated error was fixed to 5% of the average variance observed in the error free data 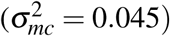.

#### 6.5.4 Simulations in Figure 4

Let *x*_*i j*_ and *y*_*i j*_ denote the intensities of the resonances measured at 3.23 and 4.98 in randomly drawn replicate *j* of sample F*i* (*i* = 1, 2, …, 5) and define the 5 × 1 vectors of means

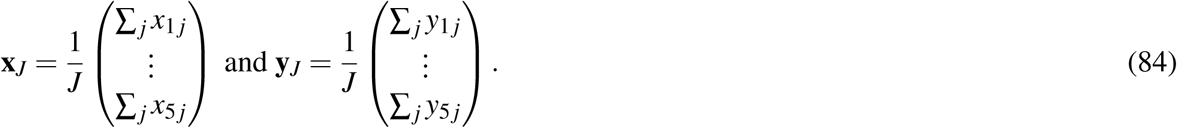

The correlation *r*_*J*_ = corr(**x**_*J*_, **y**_*J*_) is calculated for *J* = 1, 2, 5, and 10; for each *J* the replicates used to calculate **x**_*J*_ and **y**_*J*_ are randomly and independently sampled, for each sample separately, from the total set of the 12 to 15 replicates available per sample. The procedure is repeated 10^5^ times to construct the distributions of the correlation coefficent shown in Figure 4C.

#### 6.5.5 Simulations in Figure 6

Simulation results presented in Figure 6 show the results from calculations of the sample correlation coefficient as a function of the true correlation ***ρ***_0_ and of the true means (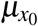 and 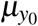), the variances (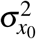 and 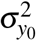 of the signals *x*_0_ and *y*_0_ and the measurement error variances as they appear in the definitions of ***ρ*** under the different error models (Equations (33), (38) and (40)). The calculations were done multiple times for varying values for 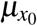 and 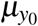, which were randomly and independently sampled from a uniform distribution *U* (0, *µ*_0_), where *µ*_0_ was set to be equal to 23.4, which was the maximum values observed in Data set 1 (see Section 6.6).Values for 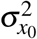 and 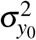 were randomly and independently sampled from a uniform distribution 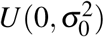, where 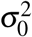 was set to be equal to the average variance observed in the experimental Data set 1. The values of the variance of all error components are randomly and independently sampled from 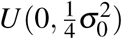. The overall procedure was repeated 10^4^ for each value of ***ρ***_0_ in the range [−1, 1] in steps of 0.1.

#### 6.5.6 Simulations in Figure 8

The first 25 variables from Data set 1 have been selected and used to compute the means *µ*_0_ and the correlation/covariance matrix Σ_0_ used to generate error-free data **X**_0_ ∼ *N*(*µ*_0_, Σ_0_) of size 10^4^ × 25 on which additive and multiplicative measurement error (correlated and uncorrelated) is added (error model (15)) to obtain **X**. All error variances are set to 0.1 which is approximately equal to 5% of the average variance observed in **X**_0_. Pairwise correlations among the 25 metabolites are calculated from **X**. The correlations are corrected using Equation (52) using the known distributional and error parameters (*µ*_0_, Σ_0_) used to generate the data. The data generation is repeated 10^3^ times and correlations (uncorrected and corrected) are averaged over the repetitions.

### 6.6 Data sets

#### Data set 1

A publicly available data set containing measurements of 133 blood metabolites from 2139 subjects was used as a base for the simulation to obtain realistic distributional and correlation patterns among measured features. The data comes from a designed case-cohort and a matched sub-cohort (controls) stratified on age and sex from the TwinGene project^44^. The first 100 observation were used in the simulation described in Section 6.5.3 and shown in Figure 3.

Data were downloaded from the Metabolights public repository^45^ (www.ebi.ac.uk/metabolights) with accession number MTBLS93. For full details on the study protocol, sample collection, chromatography, GC-MS experiments and metabolites identification and quantification see the original publication^46^ and the Metabolights accession page.

#### Data set 2

This data set was acquired in the framework of a study aiming to the “Characterization of the measurement error structure in Nuclear Magnetic Resonance (NMR) data for metabolomic studies”^29^. Five biological replicates of fish extract F1 – F5 were originally pretreated in replicates (12 to 15) and acquired using ^1^H NMR. The replicates account for variability in sample preparation and instrumental variability. For details on the sample preparation and NMR experiments we refer to the original publication.

### 6.7 Software

All calculations were performed in Matlab (version 2017a 9.2). Code to generate data under measurement error models (11), (13) and (15) is available at systemsbiology.nl under the SOFTWARE tab.

## Acknowledgements

This work was partially supported by the European Commission funded FP7 project INFECT. The authors acknowledge Peter Wentzell (Halifax, Canada) for kindly making available the NMR data set.

## Author contributions statement

E.S. and A.S conceived the study and performed theoretical calculations. E.S., M.H and A.S analysed and interpreted the results. E.S. and M.H. performed simulations. E.S., M.H and A.S wrote, reviewed and approved the manuscript in its final form.

